# A Novel *C. elegans* Memory Type Mediated by an Insulin/Phospholipase C Pathway

**DOI:** 10.1101/2023.02.22.529281

**Authors:** Daniel M. Merritt, Alexandra Udachina, Ninon Freidel, Sylvia M. T. Almeida, Yan Ming Anson Lau, Matthew Lee, Derek van der Kooy

## Abstract

Memories are often categorized into types, reflecting their behavioral, anatomical and molecular diversity: these classifications both aid understanding of the differences among varieties of memory and help delineate the unifying cross-species principles underlying them. In the nematode worm *Caenorhabditis elegans,* we find that an associative memory of the pairing of the normally attractive odorant benzaldehyde and starvation depends on *de novo* translation, is independent of CREB, and is produced by massed training: a pattern which does not correspond to any of the well-characterized molecular categories of invertebrate memory. Further, as has been shown for many memories in vertebrates, but not previously in nematodes, we find that formation of this memory continues after removal of the stimuli initially causing it, and that it is labile to disruption through protein synthesis inhibition following training, but that inhibition of proteasomal activity does not extend the duration of the memory. Previous findings have implicated insulin pathway signaling, which canonically regulates the transcription factor DAF- 16, as a key component of this benzaldehyde/starvation memory, however our results suggest that transcriptional inhibition has, at most, only moderate effects on memory formation. We find that insulin signaling instead acts to regulate phospholipase C, which in turn regulates memory through diacylglycerol signaling. These findings better characterize this model associative memory in relation to other invertebrate memory types and identify ways in which it both shares their traits and differs from them, as well as revealing a more complete picture of the molecular pathway underlying it.

## Introduction

Since at least the era of William James, researchers have understood the importance of systematically classifying the diverse phenomena of memory to understand how they function (James, 1890/2007). Memories have been categorized along many different axes, including conceptual structure (non-associative and associative memory), stimuli and paradigm (fear conditioned memory, conditioned taste avoidance memory), cognitive function (episodic memory, semantic memory, implicit memory, working memory), and by anatomical location (hippocampal memory, cerebellar memory, etc.).

Within neuroscience, one of the most influential ways of categorizing memories has been to divide them by the dissociable molecular mechanisms underlying them (and the corresponding perturbations disrupting these mechanisms). These molecular mechanisms have proven to be at least loosely correlated with the temporal sequence by which the memory develops and appear to be widely conserved, particularly in invertebrates. In the fruit fly *Drosophila melanogaster*, memories have been divided into short-term memory (STM), anesthesia-sensitive memory (ASM), anesthesia-resistant memory (ARM), and long-term memory (LTM) (Tully et al., 1994), with broadly similar divisions in *Aplysia* (Hawkins et al., 2006) (for a review of similar delineations in mammals, see Hernandez & Abel (2008)). Across all animals, most of the longest term memories require protein synthesis, as well as repeated training sessions (spaced training) rather than single training blocks (massed training) to form (Smolen et al., 2016), although there are prominent exceptions (Garcia et al., 1955).

Memory in the nematode worm *Caenorhabditis elegans* has been the subject of extensive research, and a broad assortment of genes have been identified, organized into several signaling cascades, which are required for various learning modalities. How these modalities relate to the categories of memory studied in other model invertebrates has not been clearly worked out: although types of memory are often treated as if they are broadly generalizable categories (at least within invertebrates), evidence for this remains lacking.

The role of insulin signaling in *C. elegans* memory has been extensively characterized, with many modalities depending on at least the insulin receptor DAF-2 (Cheng et al., 2022; C. H.

A. Lin et al., 2010; Tang et al., 2023; Tomioka et al., 2006). Although the *C. elegans* genome encodes approximately 40 insulin-like peptides (Hobert, 2018), benzaldehyde/starvation learning has been shown to depend solely on the ligand INS-1 (C. H. A. Lin et al., 2010). Insulin signaling in the worm canonically acts to regulate activity of the transcription factor DAF-16 (Murphy & Hu, 2013), however while DAF-16 has been shown to be required for some learning modalities (Kauffman et al., 2010; Nagashima et al., 2019; Tang et al., 2023), in others the requirement for DAF-16 is more ambiguous (Tomioka et al., 2006; Vellai et al., 2006).

To better understand how categories of memories which were primarily developed from research in fruit flies and *Aplysia* apply in *C. elegans*, we characterized a model aversive memory in which the smell of benzaldehyde, which is normally attractive to the worm, becomes temporarily aversive following paired presentation of the odor with starvation (Nuttley et al., 2002). We try to orient this memory within the categories established in the *Drosophila* literature (**Table 1**) to understand the general principles, if any, underlying molecular memory types across animals, before investigating the downstream targets of insulin signaling in this model memory.

**Table 1.** Characteristics of memory types in *Drosophila melanogaster*.

| Memory Characteristic | Memory Type |  |  |  |
| --- | --- | --- | --- | --- |
|  | Short Term Memory (STM) | Anesthesia Sensitive Memory (ASM) | Anesthesia Resistant Memory (ARM) | Long Term Memory (LTM) |
| **Protein Synthesis-Dependent** | No (Tully et al., 1994) | No (Tully et al., 1994) | No (Tully et al., 1994) | Yes (Tully et al., 1994) |
| **Spaced Training Required** | No (Davis, 2011) | No (Tully et al., 1994) | No (Tully et al., 1994) | Yes (Tully et al., 1994) |
| **Blocked by Cold Shock** | Yes (Saitoe et al., 2005) | Yes (Tully et al., 1994) | No (Tully et al., 1994) | No (Xia et al., 1998) |
| **Duration** | < 2 h (Tully et al., 1990) | ~ 7 h (Tully et al., 1994) | 2-4 days (Tully et al., 1994) | > 7 days (Tully et al., 1994) |
| **Genes Involved** | *dunce*, *rutabaga* (Tully & Quinn, 1985) | *amnesiac* (Tully et al., 1994) | *radish* (Folkers et al., 1993) | CREB (Yin et al., 1994) |

## Materials and Methods

### Strains and General Methods

All experiments were performed using wild type N2 *Caenorhabditis elegans* unless otherwise stated. Worms were grown using standard techniques at 20°C on nematode growth medium (NGM) agar plates seeded with *Escherichia coli* OP50 unless otherwise noted. All behavioral experiments were done on synchronized populations of worms 52 hours after release from L1 arrest and were performed in an environmentally controlled room at 20 °C and less than 25% humidity. Worms used in behavioral tests did not experience starvation between release from L1 arrest and initiation of training. The *C. elegans* strains N2, CF1038 *daf-16(mu86)*, RB759 *akt-1(ok525),* YT17 *crh-1(tz2)*, VC3149 *crh-2(gk3293)*, JN2722 *daf-2(pe2722)*, KQ1323 *akt-2(tm812) sgk-1(ft15)*, MT1083 *egl-8(n488)* and JN1240 *plc-1(pe1238)* were provided by the CGC, which is funded by NIH Office of Research Infrastructure Programs (P40 OD010440).

The 6x outcrossed CQ528 *pqm-1(ok485)* strain was a generous gift from Coleen Murphy. The *tpa-1(k530)* and *tpa-1(k530); pkc-1(ok563)* PKC loss of function strains were a generous gift from Anne Hart. The double CREB mutant UT1343 *crh-1(tz2); crh-2(gk3293)* was created using standard methods.

The *akt-1(mm200)* allele was generated by ethyl methanesulfonate mutagenesis (Brenner, 1974), and isolated from a mixed population following repeated selection for learning mutants in an approach similar to that of Colbert & Bargmann (1995). *mm200* contains a single base pair change resulting in an p.L199F substitution, and the originally isolated strain was outcrossed 4x to give UT1306 *akt-1(mm200)*. The *akt-1* rescue strain was created by microinjection of the fosmid WRM065aH03 into UT1306 to give UT1309.

### Transgenes and Somatic CRISPR

The constitutively active allele of *tpa-1* was designed to recapitulate the A148E mutation in the human homolog PKCθ (Liu et al., 2000), while that of *pkc-1* contained an A160E mutation in the autoinhibitory pseudosubstrate domain (Sieburth et al., 2007). Coding sequences for both genes were synthesized by Twist Bioscience, and assembled into plasmids using NEBuilder HiFi Assembly (New England Biolabs). Cell-specific expression of gain of function alleles in AWC was driven by the *ceh-36prom2* AWC promoter (Etchberger et al., 2009), while AWCoff- specific expression was driven by a promoter consisting of 2000bp upstream of *srsx-3*.

Somatic CRISPR to knockout *egl-8* specifically in chemosensory neurons was performed using a pDD162 derivative plasmid expressing a guide RNA targeting the sequence TTGATGTAGTAGTGACACAG, and Cas9 driven by a promoter consisting of 3000bp upstream of *odr-3*. Plasmids used in this work and their full sequences have been deposited with Addgene.

### Statistical Analysis

No statistical test was performed to predetermine sample sizes, which were set at 9 test plates for all experiments. The mean chemotaxis index (C.I.) and the standard error of the mean (SEM) were calculated using Excel. In most experiments with multiple independent variables, comparisons between C.I. were made by two- or three-way ANOVA (as appropriate) performed in GraphPad Prism, which were then followed by t-tests using Tukey’s multiple comparison test.

In experiments (**Fig. 2B**, **Supplemental Fig. 1**) where a full factorial design was impractical, precluding the use of a two-way ANOVA, t-tests alone were performed to test specific predicted effects, and the results adjusted for multiple comparison using Bonferroni correction. Differences were considered significant when the adjusted p<0.05. In all cases a statistical comparison of naïve chemotactic indices across matched conditions was performed, however for clarity these comparisons are only indicated on figures when a significant difference was detected. All figures depict data from three replicates performed on three different days. All statistical data are provided in **Supplemental Table 1**.

**Figure 1.**
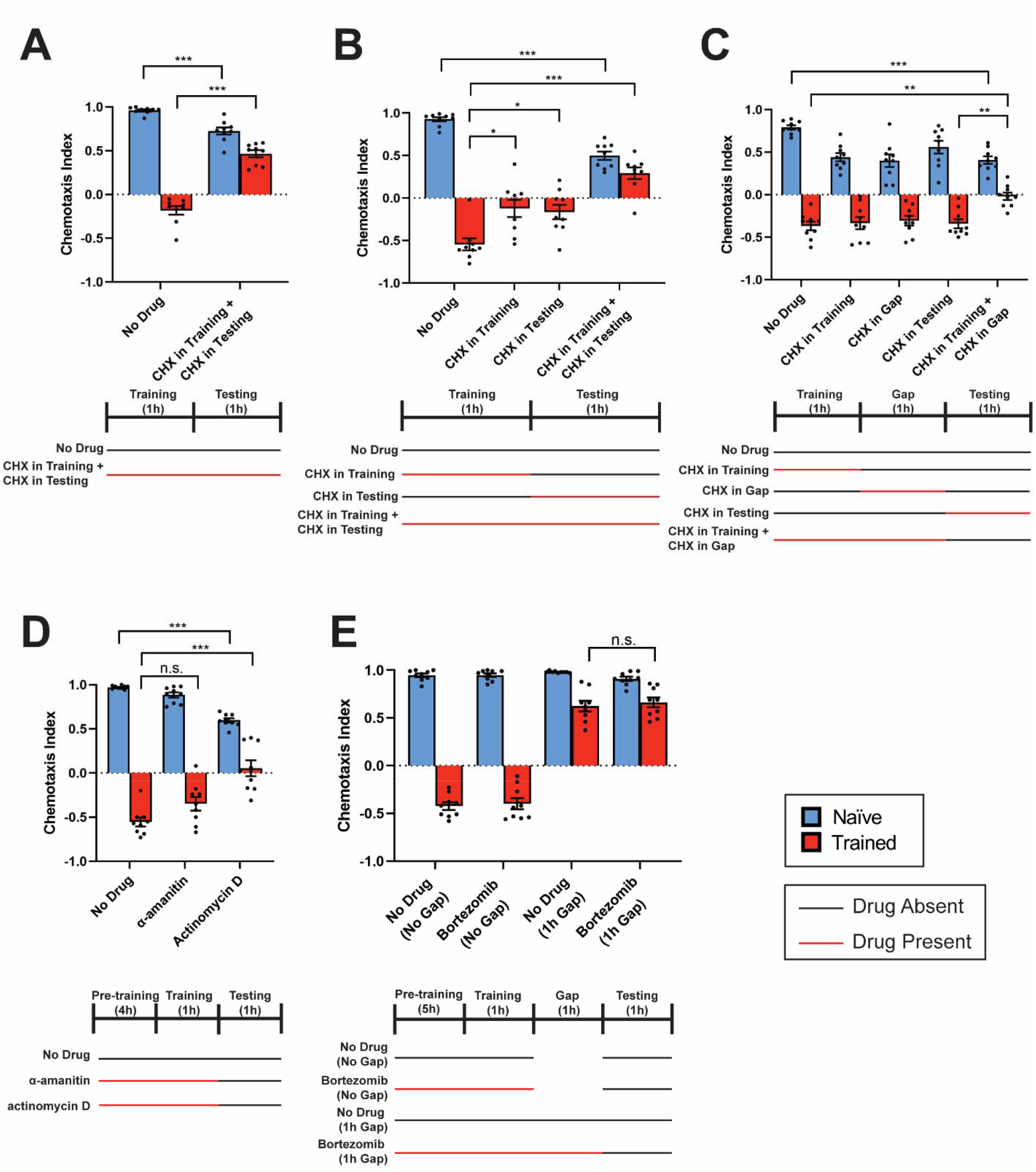
Benzaldehyde/Starvation Memory is Translation- and Transcription-Dependent. **A)** Chemotaxis of wild type animals to a point of benzaldehyde after benzaldehyde/starvation training, with and without cycloheximide in training and testing plates. **B)** Chemotaxis of wild type animals to a point of benzaldehyde after benzaldehyde/starvation training, with cycloheximide absent, present during training, present during testing, and present during training and testing. **C)** Chemotaxis of wild type animals to a point of benzaldehyde after benzaldehyde/starvation training and a 1 hour consolidation gap, with cycloheximide absent, present during training, present during the gap, present during testing or present during training and the gap. **D)** Chemotaxis of wild type animals to a point of benzaldehyde following benzaldehyde/starvation training in α-amanitin or actinomycin D, after 4h pre- exposure to the drug. **E**) Chemotaxis of wild type animals to a point of benzaldehyde after benzaldehyde/starvation training following 5 hours of proteasome inhibition by bortezomib, with and without a 1 h forgetting gap after training.

**Figure 2.**
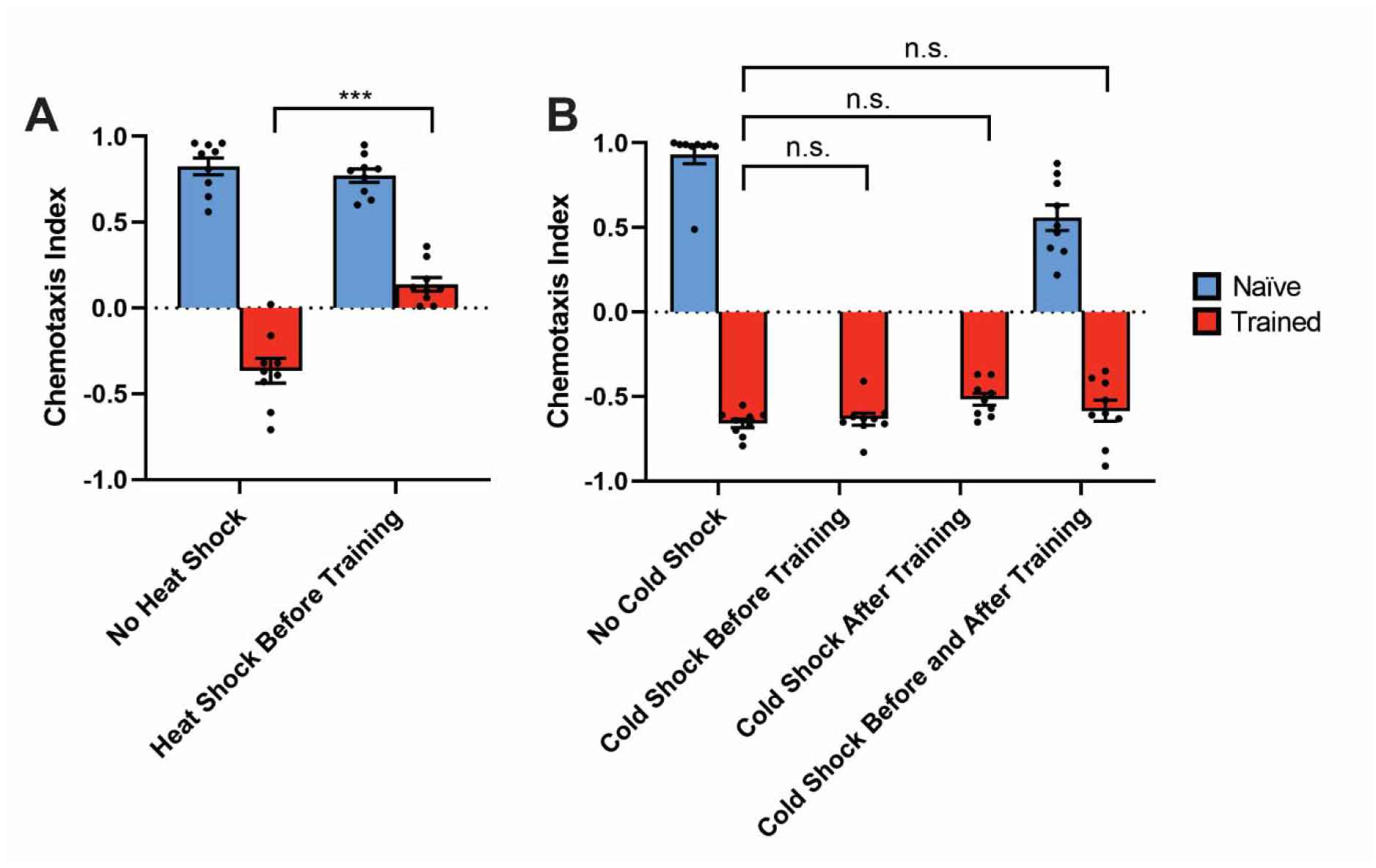
Heat Shock, but not Cold Shock, Inhibits the Benzaldehyde/Starvation Memory. **A)** Chemotaxis of wild type animals to a point of benzaldehyde after benzaldehyde/starvation training, with and without heat shock immediately before training. **B)** Chemotaxis of wild type animals to a point of benzaldehyde without cold shock, and with cold shock given before, after, and before and after training.

### CREB Phylogenetic Tree

The phylogenetic tree of CREB proteins shown in **Fig. 3A** was produced with T-Coffee sequence alignment (Notredame et al., 2000), Gblocks curation (Castresana, 2000), PhyML phylogeny reconstruction (Guindon & Gascuel, 2003) and TreeDyn rendering (Chevenet et al., 2006) using http://www.phylogeny.fr (Dereeper et al., 2008). Sequence labels were edited following tree generation for clarity.

**Figure 3.**
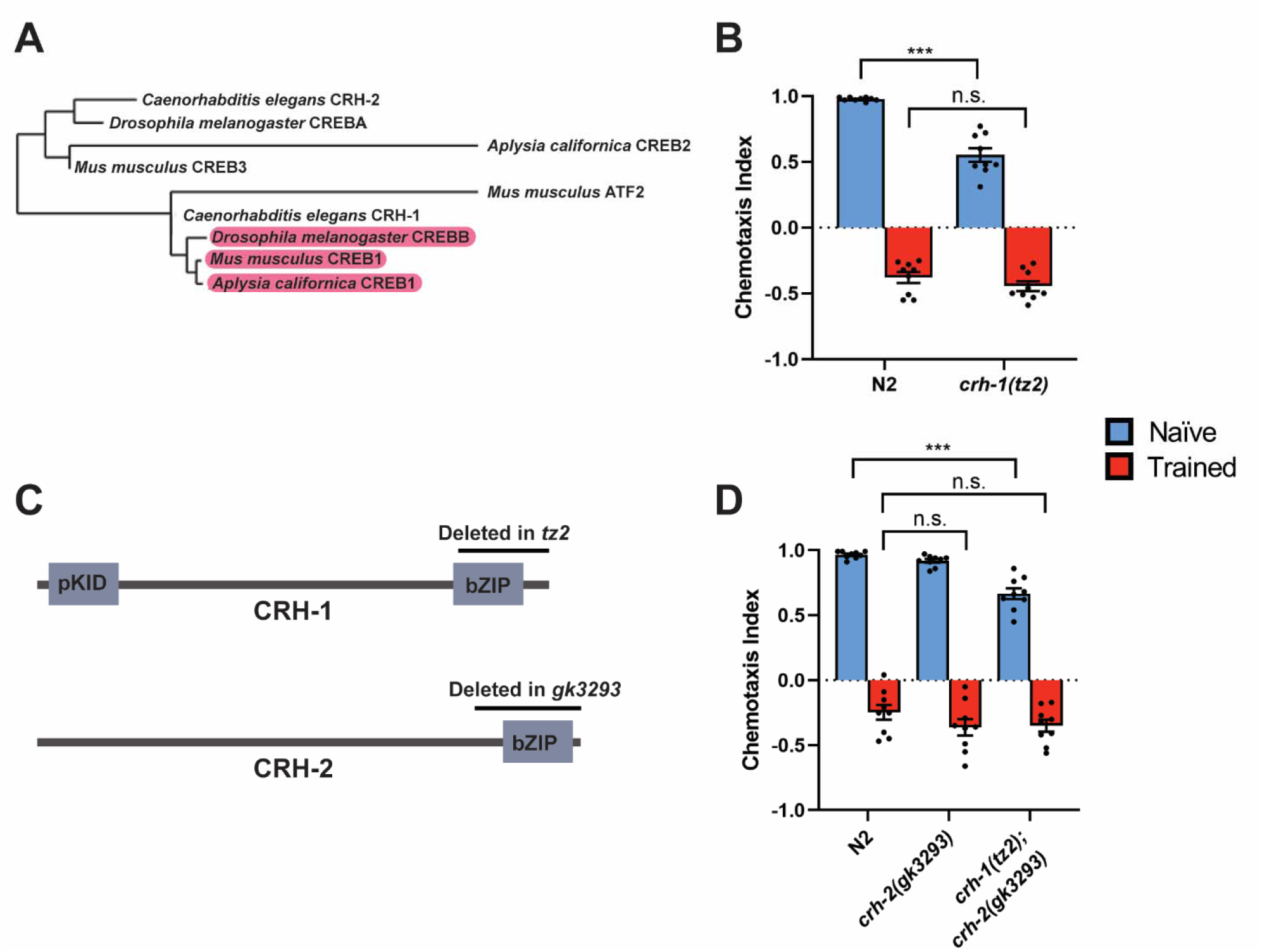
CREB is Dispensable for Benzaldehyde/Starvation Memory. **A)** Phylogenetic tree of CREB family members, with proteins implicated as positive regulators of learning highlighted in pink. **B)** Chemotaxis of wild type and *crh-1* mutant animals to a point of benzaldehyde after benzaldehyde/starvation training. **C)** Structure of *C. elegans* CRH-1 and CRH-2 proteins, with deletions tested indicated above. **D)** Chemotaxis of wild type, *crh-2* mutant and *crh-1;crh-2* double mutant animals to a point of benzaldehyde after benzaldehyde/starvation training.

Protein isoforms used for analysis were *C. elegans* CRH-1a (NP_001022859.1), CRH-2a (NP_740985.2), *Drosophila melanogaster* CREBA-PA (NP_524087.3), *Drosophila melanogaster* CREBB-PF (NP_996504.1), *Aplysia californica* CREB1 (XP_012939791.1), *Aplysia californica* CREB1 (NP_001191630.1), *Mus musculus* CREB1A (NP_598589.2), *Mus musculus* ATF-2 isoform 1 (NP_001020264.1) and *Mus musculus* CREB3 (NP_001369747.1).

### Behavioral Tests

Benzaldehyde/starvation experiments were conducted as per Nuttley et al. (2002). Briefly, learning was tested by training 1000-1500 worms for 1 hour on 6 mL, 10 cm parafilm-sealed NGM agar plates without bacteria, with either 2 µL of 100% benzaldehyde placed on a small parafilm square on the inside center of the lid (trained condition), or a parafilm square without benzaldehyde placed on the inside center of the lid (naïve condition). Following training, worms were rinsed off the plate with M9 and divided into three approximately equal groups, which were then moved to the center of three fresh 10 cm NGM testing plates, each with 1 µL of 1% benzaldehyde (Bnz) in ethanol on one side, and a second spot of 1 µL 100% ethanol (EtOH) on the other. 1 µL of 1 M sodium azide was placed on top of each odorant to paralyze the worms upon reaching it. The number of worms within 2 cm of each point of odorant, and elsewhere on the plate, was counted after 1 hour, with any worms that remained where they were placed in the center of the plate and appeared dead or injured discounted (usually < 2%), and the chemotaxis index calculated using the equation C.I.=((#Bnz)-(#EtOH))/(#Total Worms).

### Cycloheximide Treatment

Experiments utilizing cycloheximide (Bioshop) to disrupt protein synthesis were performed using 1 mL of 6 mg/mL cycloheximide dissolved in water. This was added to 5 mL of NGM agar, which had been melted and allowed to cool to below 60 °C, for a final concentration of 1 mg/mL in the training plates and/or testing plates unless otherwise stated.

For cycloheximide experiments which employed a gap between training and testing, after being washed off training plates, worms were suspended in 3 mL of M9 buffer, with or without 1 mg/mL of cycloheximide, and agitated on a rocker for 1 hour.

### α-amanitin/actinomycin D Treatment

Worms were treated with 100 µg/ml α-amanitin (Sigma) or 200 µg/ml actinomycin D (Sigma) for 4 hours prior to, and subsequently during, training. Prior to training, drugs were added to HT115 *E. coli* which had been grown overnight at 37 °C in agitated LB media before being concentrated 3 times by centrifugation followed by resuspension of the pellet in an appropriate volume of M9. During training, drugs were combined in M9 buffer along with 0.006% benzaldehyde, or M9 buffer alone for naïve controls. Tubes containing worms in buffer (+/- α-amanitin, actinomycin D, benzaldehyde or *E. coli*, as appropriate) were agitated on a rocker throughout the experiment. α-amanitin was prepared as a concentrated stock solution at 1 mg/mL in M9, while actinomycin D was dissolved at 222 µg/ml in M9.

### Bortezomib Treatment

Worms were treated with 60 µg/mL bortezomib (Cell Signaling Technology) for 5 hours prior to training, during training, and during a 1 hour gap between training and testing. Prior to training, bortezomib was added to HT115 *E. coli* which had been grown overnight (to saturation) at 37°C in agitated LB media, and then concentrated 3 times by centrifugation followed by resuspension of the pellet in an appropriate volume of M9. During training and the forgetting gap, bortezomib was added to M9 buffer along with 0.006% benzaldehyde, or M9 buffer alone, respectively. Naïve controls were treated with bortezomib in M9 buffer alone during both training and the forgetting gap. Tubes containing worms in buffer (+/- bortezomib, benzaldehyde or *E. coli*, as appropriate) were agitated on a rocker throughout the experiment. Bortezomib was prepared as a concentrated stock solution at 300 µg/mL in M9.

### Heat Shock

Worms were transferred to NGM agar plates with a lawn of *E. coli* OP50 which had been pre-warmed to 37 °C for 2 hours, and subsequently heat shocked at this temperature for 45 minutes in an air incubator immediately prior to training.

### Cold Shock

Worms were cold shocked in 3 mL ice-cold M9 buffer for 6 minutes immediately before and/or 500 µL ice-cold M9 buffer immediately after training, as indicated. Control conditions which were not cold shocked were allowed to settle in 20 °C M9 buffer for 6 minutes, before and/or after training. Training took place at 20 °C in M9 buffer containing 0.006% benzaldehyde, or in M9 buffer alone for naïve controls. Tubes containing worms in buffer (+/ benzaldehyde or *E. coli*, as appropriate) were agitated on a rocker during training.

## Results

### Benzaldehyde/Starvation Associative Memories are Protein Synthesis-Dependent

We first attempted to determine whether a model associative memory in which benzaldehyde is paired with starvation, resulting in subsequent aversion to benzaldehyde, is dependent on protein synthesis. Application of the chemical protein synthesis inhibitor cycloheximide during both initial training to avoid benzaldehyde, and during subsequent testing for benzaldehyde preference, revealed a strong effect of protein synthesis inhibition in preventing the memory, with only mild effects on naïve approach to benzaldehyde (**Fig. 1A**).

Decreasing the concentration of cycloheximide resulted in a modestly weaker learning deficit while fully eliminating the naïve approach deficit (**Supplemental Fig. 1**).

To determine when protein synthesis inhibition was acting to inhibit the memory, we applied cycloheximide selectively during training, during testing, or during both training and testing, and evaluated the strength of the resultant memory. To our surprise, cycloheximide was capable of partially inhibiting the memory during either training or testing, with the strongest inhibition resulting when it was present during both (**Fig. 1B)**. This suggested that protein synthesis was required even after removal of the training stimuli for complete learning to take place.

### Cycloheximide Inhibits Consolidation Rather than Recall

We reasoned that the inhibition of memory we observed when cycloheximide was given during testing could be due to impaired memory consolidation following training, impaired chemotaxis during testing, or impaired memory retrieval. Impaired chemotaxis was inconsistent with the larger absolute chemotaxis indices we observed in the group receiving cycloheximide during both training and testing compared to those seen in the no drug group. To distinguish between the remaining two possibilities, we added a 1 hour post-training gap between training and testing for consolidation to occur in, reasoning if the observed memory inhibition was due to impaired consolidation, cycloheximide given during this gap and during training (but not during testing) would inhibit consolidation, resulting in a memory deficit, while cycloheximide given during testing alone would have no effect. Conversely, if the memory inhibition was caused by impaired recall, cycloheximide during testing alone should retain its inhibitory effects. We find that cycloheximide during testing has no effect on memory given a 1 hour post-training gap for consolidation to occur in, suggesting that it impairs memory consolidation and that the effect of the drug during recall is in fact due to an extended period of memory formation that begins during training and continues after its end (**Fig. 1C**).

We next wondered whether benzaldehyde/starvation associative memories in *C. elegans* were dependent on *de novo* transcription, in addition to translation. We employed the transcriptional inhibitors α-amanitin and actinomycin D to inhibit transcription during training, but the results were inconclusive: while α-amanitin treatment did not result in statistically significant decreases in trained aversion, actinomycin D treatment did. The effect of actinomycin D on trained aversion was comparable to that of cycloheximide, but with a concomitant decrease in naïve approach which confounded analysis (**Fig. 1D**).

### Proteasome Inhibition does not Extend Benzaldehyde/Starvation Memory Duration

Our finding that inhibition of protein translation impairs benzaldehyde/starvation memory led us to wonder if inhibition of the proteasome might, conversely, prolong the duration of the memory by extending the lifetime of the relevant translated proteins, inhibiting an endogenous memory decay process. To test this hypothesis, we exposed worms to the small molecule proteasome inhibitor bortezomib for 5 hours prior to training, during training, and then during a 1 hour gap between training and testing: 6 hours of exposure to bortezomib at this concentration has previously been shown to be sufficient to inhibit the *C. elegans* proteasome (Melo & Ruvkun, 2012). To maximize our ability to see an increase in memory retention, we trained worms in liquid, exploiting an earlier chance observation that duration of the memory is shorter when worms are trained in liquid media than on agar plates.

We find that bortezomib-mediated inhibition of the proteasome does not result in extension of the benzaldehyde/starvation memory (**Fig. 1E)**, which is largely eliminated after 1 hour with or without bortezomib treatment, suggesting that forgetting of this memory in the worm was not mediated by the proteasome.

### Heat Shock Impairs Benzaldehyde/Starvation Memory

Inhibition of protein synthesis by heat shock has been demonstrated in yeast (Lindquist, 1981), *Drosophila* (Lindquist, 1981), mammalian cell culture (Shalgi et al., 2013) and *C. elegans* (Snutch & Baillie, 1983), and has been suggested to be mediated by pausing of translation elongation (Shalgi et al., 2013). Although not often used for classification of memories in flies and *Aplysia sp.*, heat shock has been used to impair memory in *C. elegans* (Beck & Rankin, 1995; Rose & Rankin, 2001), presumably through inhibition of protein synthesis. We find that heat shock prior to training partially blocks the benzaldehyde/starvation memory, with no effect on naïve approach to benzaldehyde (**Fig. 2A**).

### Benzaldehyde/Starvation Memory is Resistant to Cold Shock

In *Drosophila* (Quinn & Dudai, 1976), the slug *Limax flavus* (Yamada et al., 1992) and the snail *Lymnaea stagnalis* (Sangha et al., 2003), post-training cold shock is capable of disrupting the consolidation of one type of memory into another, and this effect has been suggested to be mediated by cooling-induced disruption of protein synthesis (Takahashi et al., 2013). In *Drosophila*, anesthesia-resistant memory is behaviorally differentiated from anesthesia-sensitive memory by resistance to disruption by cold shock. While some memories in *C. elegans* can be disrupted by cold shock (Morrison & van der Kooy, 1997; Nishijima & Maruyama, 2017), long-term tap habituation and long-term 1-nonanol/food appetitive learning have been reported to be cold shock-resistant (Aamodt, 2006, p. 49; Nishijima & Maruyama, 2017). To determine whether the benzaldehyde/starvation model associative memory is sensitive to cold shock, we first evaluated various durations of cold shock to determine the maximum duration we could subject worms to ice-cold M9 for without severely hindering subsequent chemotaxis, and found that 6 minutes of cold shock resulted in only a minor deficit in chemotaxis (**Supplemental Fig. 2**). This duration of cold shock is far longer than that required to disrupt memory in analogous worm paradigms (Nishijima & Maruyama, 2017), suggesting that it should be sufficient to reveal cold shock disruption if possible in our paradigm.

We previously determined that cold shock of appetitive food/salt memories in *C. elegans* had distinct effects when performed before and after training (Morrison & van der Kooy, 1997). Therefore, to determine whether this benzaldehyde/starvation memory was similarly sensitive to cold shock, we tested the effects of cold shock before, after, and both before and after training. We found that, contrary to food/salt memory but consistent with long-term tap habituation and 1- nonanol/food memory, the benzaldehyde/starvation memory was cold shock-resistant under all tested conditions (**Fig. 2B**).

### CREB is Dispensable for the Benzaldehyde/Starvation Memory

The ambiguity in our transcriptional inhibition results led us to wonder if knockout of particular transcription factors might provide more easily interpreted data on the transcriptional requirements of benzaldehyde/starvation learning. The basic region/leucine zipper (bZIP) transcription factor CREB1 has been shown to be necessary for long-term memories in diverse organisms, including *Aplysia* (Dash et al., 1990; Kaang et al., 1993), *Drosophila* (Yin et al., 1994), mice (Bourtchuladze et al., 1994) and in some paradigms *C. elegans* (Amano & Maruyama, 2011; Dahiya et al., 2019; Timbers & Rankin, 2011).

In most animals, CREB proteins exist as a diversified family, with the majority of members not having any role in memory. While the *C. elegans* genome contains two genes encoding CREB family members, *crh-1* and *crh-2,* only the protein product of *crh-1* clusters with known positive regulators of memory function in other animals (**Fig. 3A**). To determine whether the benzaldehyde/starvation memory requires CREB, we tested a strain carrying the *crh- 1(tz2)* allele. *crh-1(tz2)* mutants exhibited slightly impaired naïve approach to odorants, as has been reported previously (Dahiya et al., 2019), but we saw no evidence of a benzaldehyde/starvation learning deficit (**Fig. 3B**), consistent with our previous observations on these mutants in more complex memory paradigms (Merritt et al., 2019). Since the *crh-1(tz2)* allele deletes most of the bZIP domain of *C. elegans* CREB1 **(Fig. 3C)**, and the allele results in the elimination of CREB as detected by anti-phospho-CREB antibody (Kimura et al., 2002), it is a presumptive null, and we therefore conclude that the benzaldehyde/starvation memory is CRH- 1-independent.

While homology and behavioral evidence suggested that CRH-1 was the most likely CREB family candidate involved in this memory, it remained possible that the CRH-2 was the relevant CREB protein. We therefore tested a strain carrying the *crh-2(gk3293)* allele, which eliminates the entire CRH-2 bZIP domain and is therefore a probable null, but found no evidence of a learning deficit. A double mutant strain carrying mutations in both *crh-1* and *crh-2* shows no additive phenotype over the *crh-1* mutant strain, suggesting that our failure to observe a learning deficit was not due to genetic redundancy **(Fig. 3D)**.

### The Insulin Signaling Pathway is Required for Benzaldehyde/Starvation Learning at Least as far as AKT-1

Previous work from our group and others (Cheng et al., 2022; C. H. A. Lin et al., 2010; Tomioka et al., 2006) has described a key role for insulin signaling in olfactory and gustatory learning in *C. elegans*. The extensive transcriptional changes known to take place downstream of insulin signaling (Murphy, 2006) suggested a straightforward explanation: could the requirement we observed for de novo translation (and potential requirement for transcription) in the benzaldehyde/starvation learning paradigm result from insulin signaling-mediated changes in gene expression?

Canonical regulation of transcription through the insulin signaling pathway is mediated by two downstream transcription factors with opposing regulatory effects: DAF-16 (Schuster et al., 2010) and PQM-1 (Tepper et al., 2013). DAF-16 is excluded from the nucleus by phosphorylation by the kinase AKT-1 when the insulin signaling pathway is active (Paradis & Ruvkun, 1998), but in the absence of upstream signaling enters the nucleus to transcriptionally activate target genes (class I genes). PQM-1 is excluded from the nucleus by nuclear DAF-16, and so only in the presence of insulin signaling is it able to enter the nucleus to transcriptionally activate its targets (class II genes) (Tepper et al., 2013).

We first sought to clarify whether the entire canonical insulin signaling pathway was required for benzaldehyde/starvation associative learning. Previous work has shown a requirement for the sole insulin receptor *daf-2*, its ligand *ins-1* and the downstream kinase *age-1* in this paradigm (C. H. A. Lin et al., 2010), however since then, the requirement for DAF-2 in other *C. elegans* learning modalities has been shown to be specific to the DAF-2c isoform (Tomioka et al., 2016, 2022). We first tested a *daf-2c* isoform-specific loss of function mutant, and determined that it fully blocks learning in our paradigm (**Fig. 4A**). Since in the canonical insulin signaling pathway, the final step before transcription factor involvement is phosphorylation of the serine/threonine kinase AKT-1, we next tested mutants in *akt-1* to confirm the necessity of the pathway up to the point of transcriptional regulation. We find that *akt-1* mutant animals exhibit a severe abrogation of learning (**Fig. 4B**) which can be rescued by wild type *akt-1* (**Supplemental Fig. 3**). This confirmed that the requirement for the insulin signaling pathway extends at least as far in the pathway as *akt-1*.

**Figure 4.**
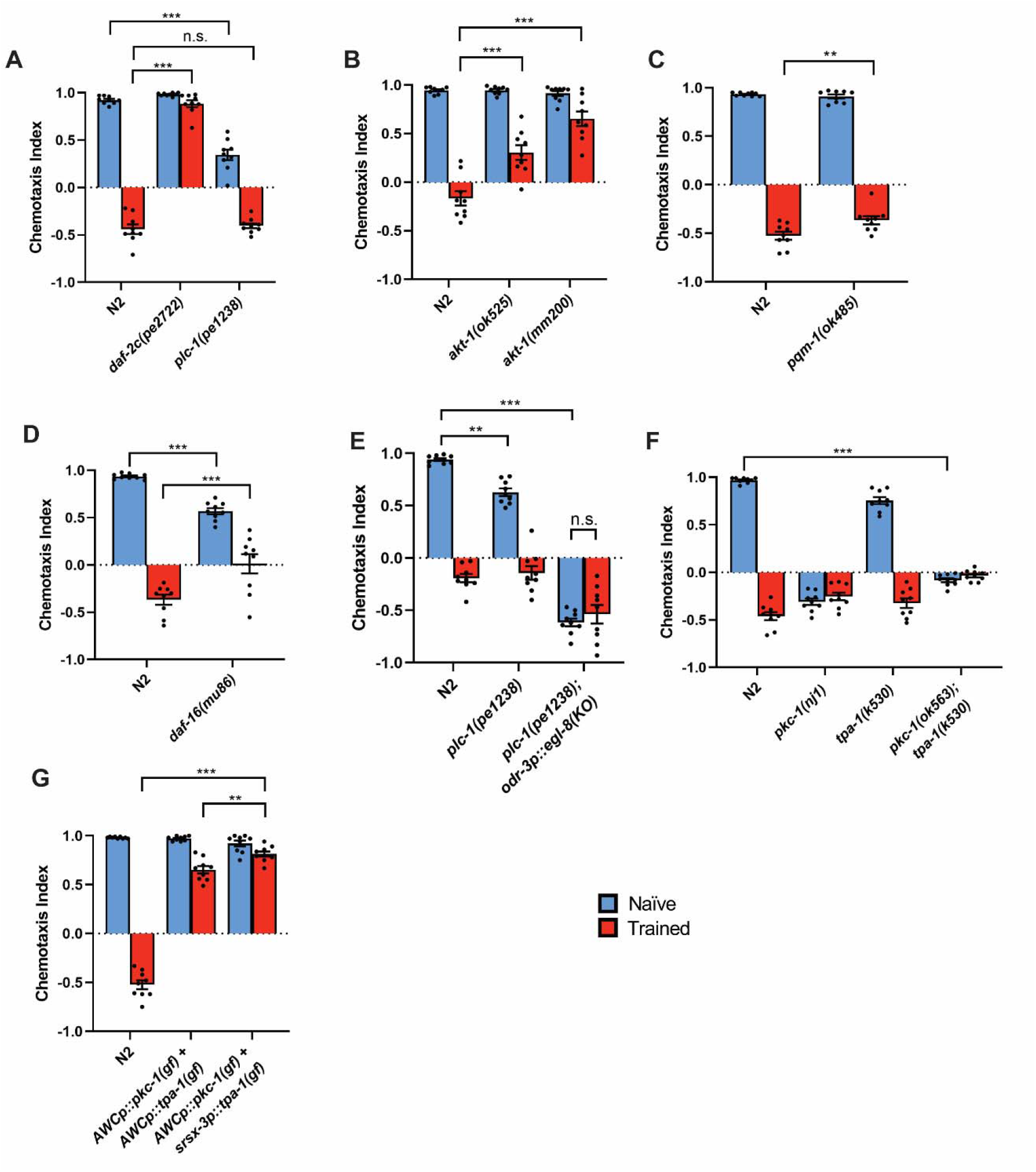
Benzaldehyde/Starvation Learning Depends on Insulin Signaling but not Canonical Insulin Signaling Transcription Factors. **A)** Chemotaxis of wild type, *daf-2c* and *plc-1* mutant animals to a point of benzaldehyde after benzaldehyde/starvation training. **B)** Chemotaxis of wild type and *akt-1* mutant animals to a point of benzaldehyde after benzaldehyde/starvation training. **C)** Chemotaxis of wild type and *pqm-1* mutant animals to a point of benzaldehyde after benzaldehyde/starvation training. **D)** Chemotaxis of wild type and *daf-16* mutant animals to a point of benzaldehyde after benzaldehyde/starvation training. **E)** Chemotaxis of wild type and *plc-1* mutant animals, and *plc-1* mutant animals with *egl-8* selectively knocked out in chemosensory neurons, to a point of benzaldehyde after benzaldehyde/starvation training. **F)** Chemotaxis of wild type and *pkc-*1, *tpa-*1 mutants, and *pkc-1;tpa-1* double mutant animals, to a point of benzaldehyde after benzaldehyde/starvation training. **G)** Chemotaxis of wild type animals, animals expressing constitutively active alleles of *pkc-1* and *tpa-1* expressed in both AWCs, and animals expressing a constitutively active allele of *pkc-1* in both AWCs, in addition to a constitutively active allele of *tpa-1* expressed in AWCoff, to a point of benzaldehyde after benzaldehyde/starvation training.

Since AKT-1 activity results in the nuclear localization of PQM-1 (via nuclear exclusion of DAF-16), we reasoned that PQM-1 was the downstream transcription factor most consistent with our observed effects of transcriptional inhibition. We therefore tested *pqm-1* loss of function mutant animals (Tepper et al., 2013) for learning deficits, but to our surprise found that this strain exhibited only an extremely weak learning deficit (**Fig. 4C**). *daf-16* null animals (K. Lin et al., 1997) were also capable of learning (**Fig. 4D**), however they showed a mild deficit in naïve approach to benzaldehyde similar to that previously reported in this strain (Tomioka et al., 2006). This impaired approach made it difficult to conclusively rule in or out an impairment of learning. Since AKT-1 is a negative regulator of DAF-16, it might be expected that a *daf-16* mutant would exhibit a “constitutively trained” phenotype if it was the sole substrate of AKT-1 in benzaldehyde/starvation learning, however we saw only mildly impaired approach to benzaldehyde (which was equally consistent with impaired movement or chemosensation) and continued to see aversion only after training, inconsistent with this hypothesis.

Since we found that *akt-1* loss of function mutations do not totally block learning, we wondered whether *akt-2* or *sgk-1*, which operate in parallel to *akt-1* for some phenotypes regulated by insulin signaling (Chen et al., 2013; Paradis & Ruvkun, 1998), might mediate the small degree of behavioral learning remaining in *akt-1* mutants in our assays. *akt-1;akt-2* double mutants are constitutively dauer (Paradis & Ruvkun, 1998), preventing us from studying this double (or the triple) mutant, so we tested an *sgk-1;akt-2* double mutant to see if it partially abrogated learning. We found that, contrary to the prediction of this model, this double mutant exhibited slightly improved learning (**Supplemental Fig. 4**), consistent with a prior report of *akt-2* being uninvolved in salt learning (Tomioka et al., 2022) and perhaps consistent with the reported role of *sgk-1* acting opposite *akt-1* in regulating lifespan (Chen et al., 2013).

### AKT-1 Negatively Regulates Phospholipase C to Mediate Learning

The failure of *daf-16* mutants to exhibit a constitutively learned phenotype, and the retention of experience-dependent aversion in these mutants, suggested that DAF-16 was not the sole downstream target of AKT-1 in benzaldehyde/starvation learning. Prior work has identified AKT-1 phosphorylation motifs on the phospholipase C (PLC) enzyme PLC-1, and identified a role for PLC-1 and a second PLC, EGL-8, downstream of AKT-1 (Tomioka et al., 2022).

Consistent with this, we also found an AKT-1 phosphorylation motif (Alessi et al., 1996) in EGL-8 (but not in the homologous PLCs *plc-2*, *plc-3*, *plc-4* or *pll-1*) (**Supplemental Fig. 5**). PLCs canonically act to cleave phosphatidylinositol 4,5-bisphosphate into diacylglycerol (DAG) and inositol 1,4,5-trisphosphate, each of which result in distinct downstream signaling cascades (Vines, 2012), and DAG has previously been strongly implicated in *C. elegans* chemotaxis and olfactory learning (C. H. A. Lin et al., 2010; Matsuki et al., 2006; O’Halloran et al., 2009; Tsunozaki et al., 2008).

Loss of function mutations in each of *egl-8* and *plc-1* resulted in naïve approach deficits which confounded analysis. RNAi against *egl-8* has been reported to result in slow growth, which we also observed in our *egl-8* mutant strain, and we wondered if the observed naïve approach deficit could result from a failure to reach young adulthood in this strain. To compensate for this, we grew *egl-8* mutant animals at a warmer temperature, which allowed worms to reach adulthood under our normal assay’s 52 hour development time, but this did not restore strong naïve approach (**Supplemental Fig. 6**).

An alternative explanation was suggested by the prior findings that in high salt starvation training conditions, EGL-8 and PLC-1 are negatively regulated by AKT-1 to mediate movement toward low salt (Tomioka et al., 2022), and that loss of function mutations in these PLCs could reduce benzaldehyde attraction (O’Halloran et al., 2009). This led us to wonder if a similar mechanism could underlie trained aversion to benzaldehyde in our paradigm: rather than being reflective of movement or chemosensory disorders, the poor naïve approach seen in the naïve condition in *egl-8* and *plc-1* mutant animals might result from a genetic recapitulation of AKT-1- mediated downregulation of EGL-8 and PLC-1, resulting in a constitutively trained animal.

Since previous evidence has implicated the chemosensory neuron AWC as the required site of *age-1* activity during benzaldehyde/starvation learning (C. H. A. Lin et al., 2010), we used somatic CRISPR (Shen et al., 2014) to knock out *egl-8* selectively in chemosensory neurons in a *plc-1* mutant background. Consistent with this hypothesis, we found that animals exhibited robust chemotaxis away from benzaldehyde in both trained and naïve conditions (**Fig. 4E**).

### Protein Kinase C Acts Downstream of Diacylglycerol to Promote Benzaldehyde Approach

Members of the Protein Kinase C family are common upstream effectors of DAG signaling, including in some learning modalities (Hiroki & Iino, 2022; Stetak et al., 2024). Since previous reports suggested that loss of function mutations in the PKC family member *pkc-1* resulted in strong aversion to the AWCon-sensed odorant butanone, but only minor chemotaxis deficits to the AWCon and AWCoff-sensed odorant benzaldehyde (Tsunozaki et al., 2008), we wondered if a second PKC might act redundantly with PKC-1 in AWCoff to mediate behavioral effects downstream of DAG. Since single-cell RNA sequencing suggested that the PKC TPA-1 was more strongly expressed in AWCoff than AWCon (Taylor et al., 2021), and since some previous reports had suggested a minor role for it in benzaldehyde chemotaxis (Okochi et al., 2005), we considered it as a candidate.

We first examined loss of function mutants of *tpa-1* and *pkc-1* (Okochi et al., 2005). Loss of function mutations in *pkc-1* resulted in modest constitutive benzaldehyde avoidance in our paradigm, consistent with previous reports (Okochi et al., 2005), while a *tpa-1* mutation resulted in slightly diminished approach, consistent with *pkc-1* playing a primary role. A double mutant strain exhibited severely abrogated chemotaxis, suggesting either a loss of chemosensation or broader pleiotropic effects (**Fig. 4F**). These results were consistent with PKC-1 playing the major role in DAG-mediated aversion, but did not rule out a minor role for other PKCs, including TPA-1.

If DAG production by PLCs is negatively regulated by AKT-1 during learning, and this mediates trained benzaldehyde aversion, constitutive activation of PLCs would be expected to block benzaldehyde learned aversion. To test this prediction, we constructed a strain containing predicted gain of function alleles of *pkc-1* and *tpa-1* expressed under an AWC-specific promoter, and found that, as our model predicts, this strain exhibited a nearly complete block of learning.

Since previous behavioral data suggests that *tpa-1* activity may be specific to AWCoff, and single cell RNA sequencing suggested weaker expression in AWCoff relative to AWCon (Taylor et al., 2021), we also constructed a strain in which a constitutively active allele of TPA-1 was selectively expressed in AWCoff in addition to dual AWC expression of a constitutively active allele of PKC-1. We observed a robust but incomplete block of learning in both strains, consistent with our model (**Fig. 4G**). Residual learning in these strains is likely caused by mosaicism of the extrachromosomal array carrying the constitutive allele.

## Discussion

### Protein Synthesis-Dependent Consolidation Continues After Training

Our data suggest that withdrawal of protein synthesis inhibition after completion of training allows for partial formation of the benzaldehyde/starvation memory. When training is immediately followed by testing, inhibition of protein synthesis during either phase results in significant abrogation of memory, and inhibition of protein synthesis during both results in nearly total loss of memory. When a gap period is interspersed between training and testing phases, however, preventing protein synthesis during testing or training individually no longer impairs memory: memory formation is only blocked, and even then not fully, when protein synthesis is inhibited during both training and the gap period. This pattern of results suggests that there is a wide temporal window for protein synthesis-dependent memory formation to occur in, and further, that there is no necessary requirement for protein synthesis during memory recall.

Post-training consolidation of memory has been extensively studied in mammals, flies and *Aplysia*, primarily in the context of long-term memories. Consolidation in these organisms can be disrupted by protein synthesis inhibition, with a critical period of a few hours post-training generally constituting the window during which protein synthesis inhibitors are effective (Flexner et al., 1965; Goelet et al., 1986; Wu et al., 2017) except during episodes of subsequent reconsolidation (Nader et al., 2000). Our data support a similar mechanism in formation of the benzaldehyde/starvation memory, whereby external stimuli set in motion memory formation processes which continue after their removal.

### The Identity of Gene Products Requiring Translation Remains Unclear

Many genes have been implicated in *C. elegans* memory, and in principle translation of any of them might be required during learning. Our chemical inhibition data suggest that *de novo* translation of previously inactive genes is required during learning. This constrains the range of possible genes to those which are not normally translated in the relevant cell type, and to protein coding genes specifically (ruling out, for example, *odr-1-*targetting endo-siRNAs (Juang et al., 2013)).

The key role played by insulin signaling during benzaldehyde/starvation associative learning, and the well-studied termination of this pathway in DAF-16- and PQM-1-mediated transcriptional changes, make these an appealing mechanism to explain our drug inhibition effects, however the data we present here suggest it is unlikely to be the causative mechanism. Our present findings, in conjunction with the previous identification of *ins-1*, *daf-2* and *age-1* as required genes (C. H. A. Lin et al., 2010), support the necessity of components of the insulin signaling pathway as far downstream as AKT-1, but *pqm-1* mutants exhibit only extremely small learning deficits relative to those caused by translational inhibition. *daf-16* mutant animals are more ambiguous because of a substantial naïve approach deficit, however the learning deficit present in the trained group is still weaker than that caused by protein synthesis inhibition, suggesting cycloheximide cannot be acting by inhibiting translation of DAF-16-mediated transcriptional targets. A requirement for *daf-16-*mediated transcription is also mechanistically inconsistent with a requirement for *akt-1*, since loss of function *akt-1* mutations result in the constitutive nuclear localization of DAF-16, but result in abrogation of learning. These findings suggest that the role of the insulin signaling pathway in learning is not mediated by its canonical downstream transcription factors DAF-16 and PQM-1. Evidence for DAF-16-independent roles of the insulin signaling pathway in learning has previously been identified in the worm (Tomioka et al., 2006), and highly paradigm-specific roles for various components of the pathway have been suggested (Nagashima et al., 2019).

Research in other learning modalities in *C. elegans* has shown that regulation of experience-dependent plasticity in synaptic size, thought to be a mechanism of learning, requires translation of the ARP2/3 complex, and that this is negatively regulated by the pro-forgetting protein MSI-1 (Hadziselimovic et al., 2014). This represents a second potential mechanism by which translational inhibitors may impair memory in *C. elegans*.

Our finding that inhibition of the proteasome does not extend memory duration is consistent with both a mechanism in which protein synthesis is necessary for formation of the memory engram but does not itself constitute it, or with a proteasome-independent mechanism of engram decay (for example, free cellular proteases or autophagy). Genes mediating active processes of forgetting in *C. elegans* have been identified (Arai et al., 2022; Bai et al., 2022; Hadziselimovic et al., 2014; Inoue et al., 2013; Teo et al., 2022), however none have encoded proteins involved in protein decay, consistent with our findings.

### Benzaldehyde/Starvation Memory Exhibits Characteristics of Several Invertebrate Memory Types

In fruit flies and *Aplysia*, memories produced by a single training session (massed training) are generally independent of transcription, translation and the transcription factor CREB, while repeated training sessions (spaced training) give rise to long-term memories dependent on transcription, translation and CREB (**Table 1**). The *C. elegans* memory we describe here does not clearly correspond to any of these: it depends on transcription and translation, is independent of CREB, and is produced through a single massed training session.

Two very different interpretations of these results can be imagined. It could be that the memory described here is a genuinely independent class of memory with different behavioral and molecular requirements to those of other well-studied invertebrate memories. Alternatively, it may be that the behavioral manifestation of this memory is mediated by an overlapping suite of memory classes which independently correspond to the memory types seen in flies and *Aplysia*, but which in conjunction manifest as a blended memory mechanism. However, since this latter interpretation would at the very least require postulating a protein synthesis-dependent memory produced by massed training as one of these constituent components, the requirement for a novel class of memory cannot be fully avoided, and so we favor the former interpretation as more parsimonious.

Massed training protocols in the worm have been reported to be either independent (Amano & Maruyama, 2011; Nishijima & Maruyama, 2017) or dependent (Stein & Murphy, 2014) on *de novo* protein translation, depending on the protocol. While it is conceptually possible that some apparently massed protocols involve cryptic spaced training, it is difficult to see ways by which this could occur in ours: food is uniformly absent on the agar plate (or in liquid media) during training, and the worms are exposed to the smell through the gas phase, which is ubiquitously present. Our findings, in conjunction with a similar report of protein synthesis-dependent massed learning using a different paradigm (Stein & Murphy, 2014), challenge the view that protein synthesis-dependent memory necessarily requires spaced training: in *C. elegans,* requirement for spaced training appears instead to depend on the exact conditioned and/or unconditioned stimuli employed.

None of the chemical manipulations we describe here fully prevent memory. It is possible that the small fraction of retained memory in our experiments represents a (potentially short-term) memory process which is independent of translation and transcription. Since experiments in a similar paradigm found that only 50% of protein synthesis was inhibited by a concentration of cycloheximide three times greater than the one we use (Amano & Maruyama, 2011), however, at least for our experiments utilizing cycloheximide, limited drug efficacy is a more likely explanation.

### A Molecular Model of Benzaldehyde/Starvation Memory

The molecular analysis of memory formation here suggests a model of part of the genetic pathway by which benzaldehyde/starvation memory is produced. We propose a model in which insulin signaling acts on DAF-2c in the AWCs, leading to activation of AKT-1, which in turn negatively regulates the PLCs EGL-8 and PLC-1 by phosphorylation, resulting in learned aversion to benzaldehyde (Fig. 5). Conversely, in an unphosphorylated state, these PLCs produce PIP-3 and DAG, the latter of which activates PKC-1 and possibly TPA-1, leading to positive benzaldehyde chemotaxis. Aspects of this model are supported by results recently described in a preprint by another group (Berliner et al., 2024).

**Figure 5.**
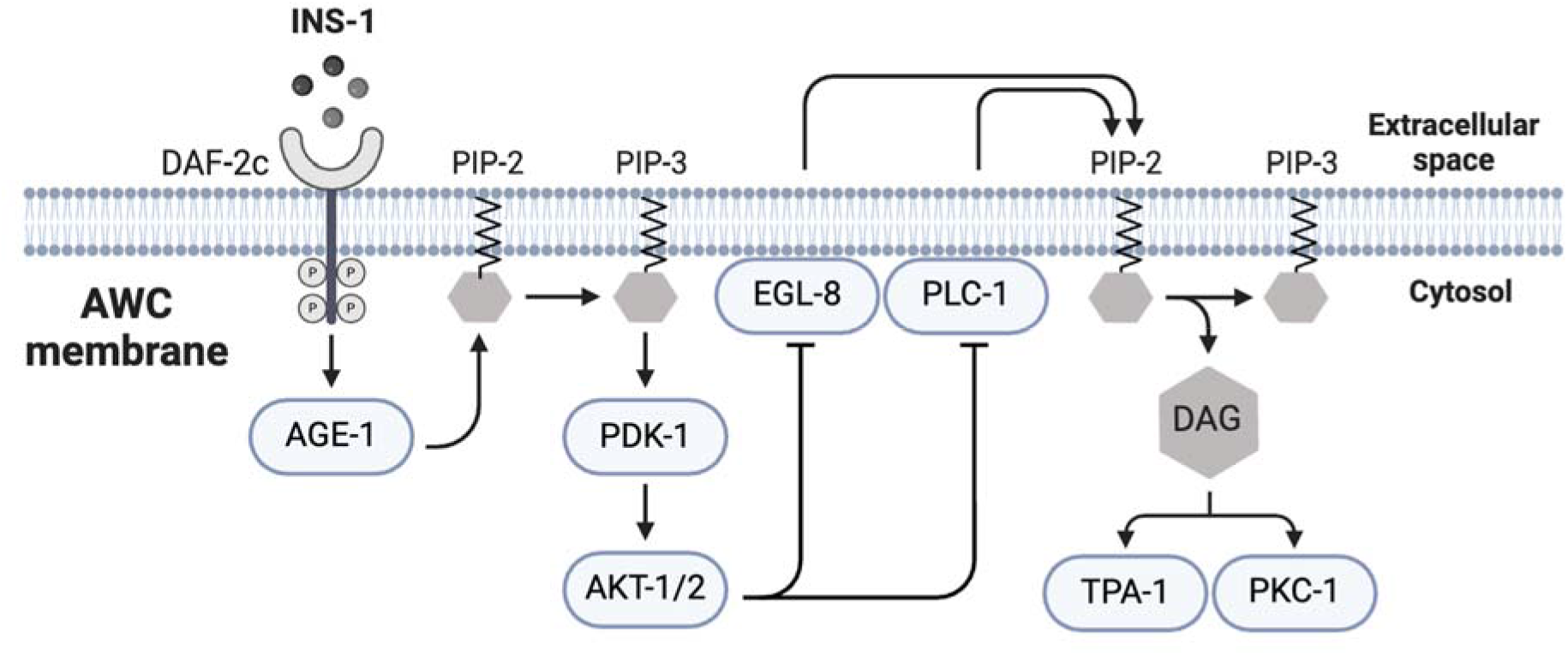
Proposed Partial Model of Benzaldehyde/Starvation Memory Pathway in AWC. We propose a model in which benzaldehyde/starvation learning proceeds via INS-1, acting on the DAF-2c receptor, which activates the insulin signaling cascade to phosphorylate AKT-1 and potentially AKT-2. Phosphorylated AKTs then negatively regulate the PLCs EGL-8 and PLC-1, inhibiting production of DAG. DAG normally stimulates the PKCs PKC-1 and potentially TPA-1, leading to benzaldehyde attraction, and decreased DAG levels result in benzaldehyde aversion. Grey hexagons represent small molecules, while blue ovals represent proteins.

In *egl-8* isoforms which contain it, the AKT binding motif is present in a very short exon consisting of 12 amino acids, 7 of which constitute this binding motif. We speculate that this may constitute a mechanism by which insulin signaling can be linked to PLC signaling by an alternative splicing event, perhaps determined by factors such as cellular or environmental context.

Several aspects of this model remain unclear. Loss of function mutations in *akt-1* block most, but not all, benzaldehyde/starvation learning, while *daf-2c* and *ins-1* mutations result in a full block of learning. Residual learning in *akt-1* mutants may be mediated by redundant activity of *akt-2*. While our data suggest that DAG primarily acts through negative regulation of PKC-1 in learning, a minor role for TPA-1 remains possible. As well, the targets downstream of these PKCs remain unclear: PKCs have many downstream effectors, and which ones are relevant for our learning modality have yet to be determined.

The molecular pathway described here both shares similarities and exhibits differences from those identified for other learning modalities in *C. elegans*. Butanone learning, in which the primary sensory neuron is AWCon, might be expected to share a broadly overlapping set of molecular components to benzaldehyde/starvation learning, where the olfactory cue is sensed by AWCon and AWCoff. The failure of these stimuli to cross-adapt, however, suggests that at least some differences must exist (Colbert & Bargmann, 1995).

Learning from your mistakes is a valuable skill for all animals, and for most animals there are many different mistakes which might profitably be learned from. Our findings suggest that memory types across invertebrates, and even different memories within a single species, may exhibit fewer unifying principles than is commonly appreciated, at least as revealed by the systems of classification which have been influential in flies and *Aplysia*. This both highlights the importance of considering the specific paradigm and species in question when reasoning about the mechanisms of memory processes, and draws attention to the diversity of ways in which evolution has provided mechanisms to change behavior in response to a dangerous world.

## Acknowledgements

This research was supported by Natural Sciences and Engineering Council of Canada (NSERC). Some strains were provided by the CGC, which is funded by NIH Office of Research Infrastructure Programs (P40 OD010440). We thank Coleen Murphy for the generous gift of the 6x outcrossed *pqm-1* strain, and Anne Hart for the *tpa-1(k530)* single and *tpa-1(k530); pkc-1(ok563)* double mutant PKC strains. We thank Sebastien Santini (CNRS/AMU IGS UMR7256) and the PACA Bioinfo platform for the availability and management of the phylogeny.fr website used for the analysis depicted in Fig. 3A. Fig. 5 and **Supplemental** Fig. 5 were created with BioRender.com.

## Author contributions

D.M.M.: Conceptualization, Data Curation, Formal Analysis, Project Administration, Supervision, Visualization, Writing – Original Draft Preparation, Writing – Review and Editing.

A.U.: Data Curation, Formal Analysis, Investigation, Methodology, Conceptualization, Writing – Review and Editing.

N.F.: Data Curation, Investigation, Conceptualization, Visualization, Writing – Review and Editing.

S.M.T.A.: Formal Analysis, Investigation, Conceptualization, Writing – Review and Editing.

Y.M.A.L.: Investigation, Validation, Writing – Review and Editing. M.L.: Investigation, Validation, Writing – Review and Editing.

D.v.d.K.: Conceptualization, Project Administration, Supervision, Funding Acquisition, Writing – Review and Editing.

## Data and materials availability

All data are available in the main text or the supplementary materials.

