## Supplemental Figures for "A Novel *C. elegans* Memory Type Mediated by an Insulin/Phospholipase C Pathway"

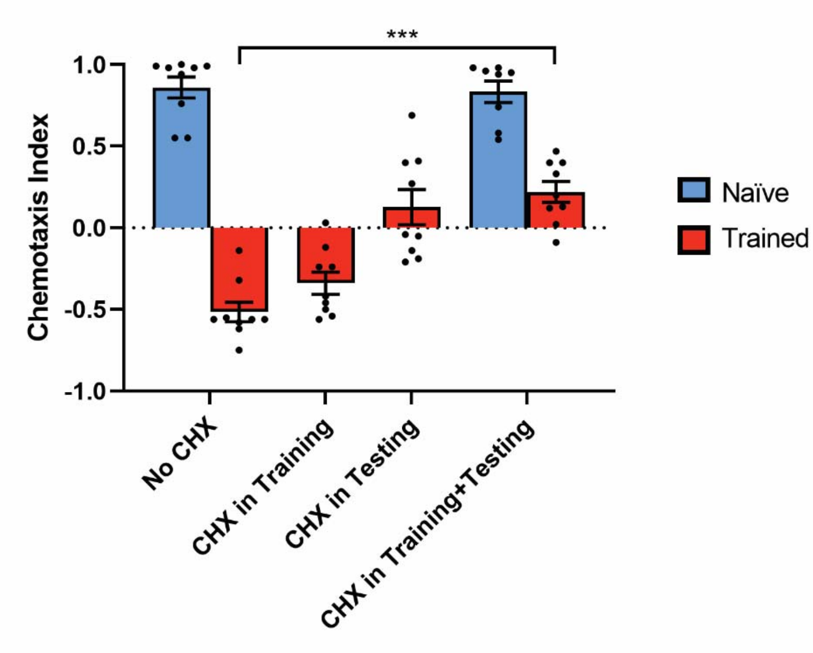


**Supplemental Figure 1** Protein Inhibition by a Lower Concentration of Cycloheximide Inhibits Memory without Impairing Movement. Chemotaxis of wild type animals to a point of benzaldehyde after benzaldehyde/starvation training, with cycloheximide absent, present at a concentration of 0.25mg/mL during training, during testing, and during training and testing.

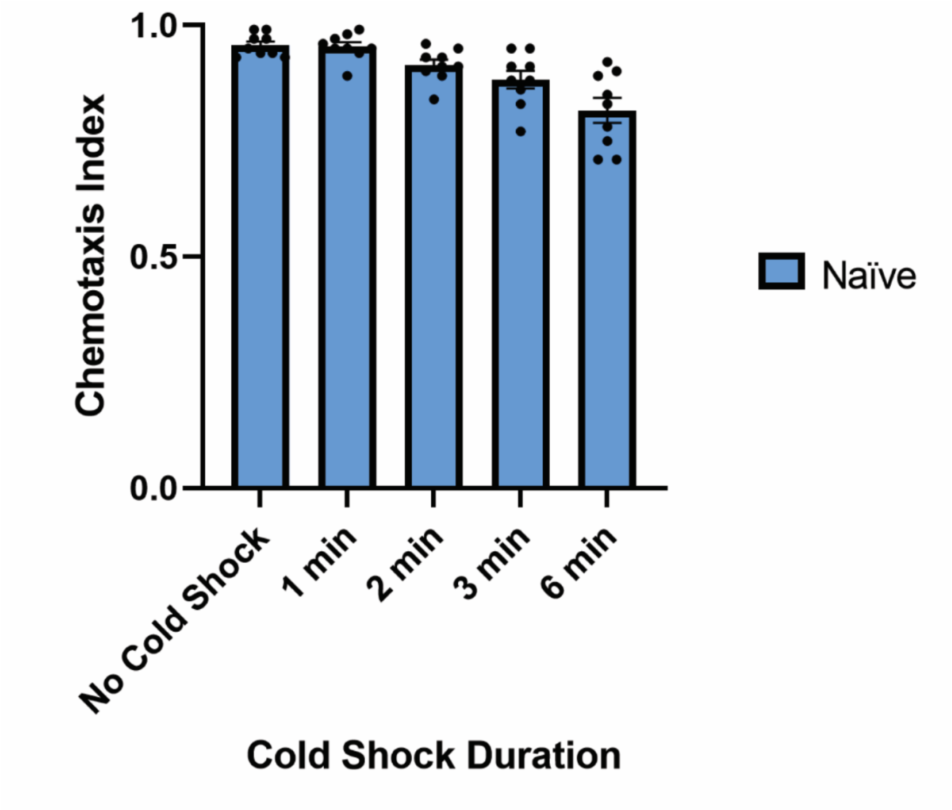


**Supplemental Figure 2** Six Minutes of Cold Shock Results in Only Minor Chemotactic Deficits. Chemotaxis of wild type animals to 1% benzaldehyde after 1 hour starvation, immediately following cold shock.

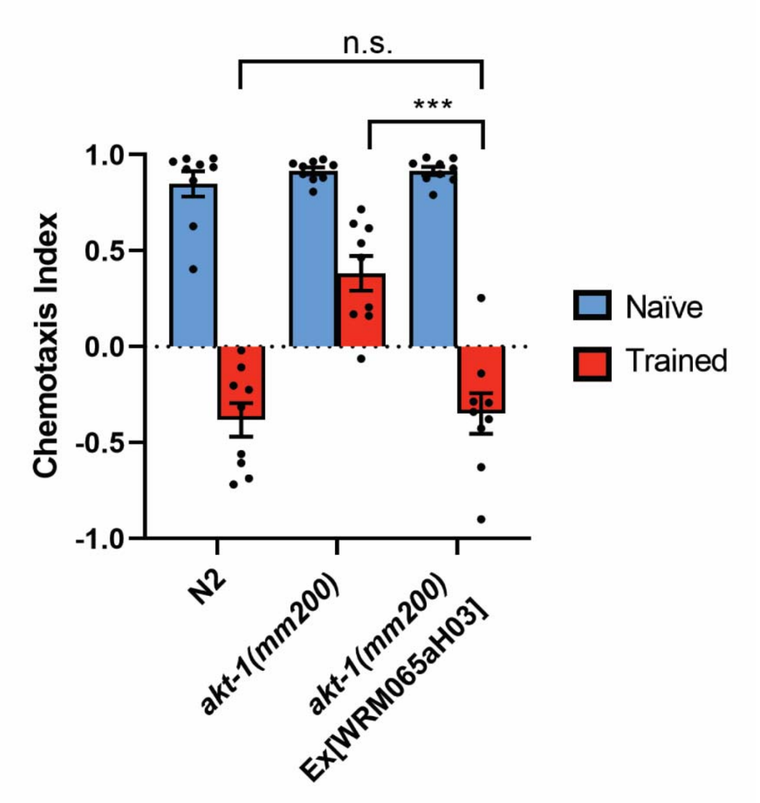


**Supplemental Figure 3** A fosmid containing wild type *akt-1* restores learning when carried as an extrachromosomal array. Chemotaxis of wild type, *akt-1(mm200)* and *akt-1(mm200)* worms carrying the fosmid WRM065aH03 (which contains wild type *akt-1*) as an extrachromosomal array, to a point of benzaldehyde after benzaldehyde/starvation training.

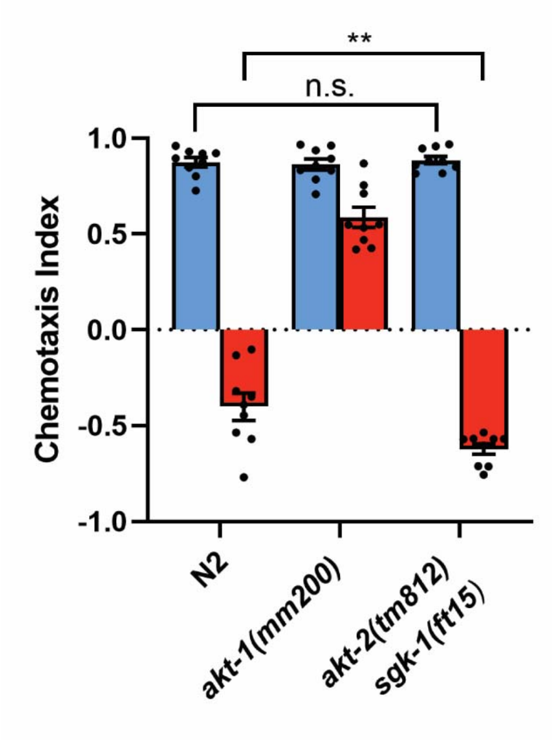


**Supplemental Figure 4** Chemotaxis of N2 animals, *akt-1(mm200)* animals and *akt-2(tm812) sgk-1(ft15)* double mutant animals to a point of benzaldehyde after benzaldehyde/starvation training.

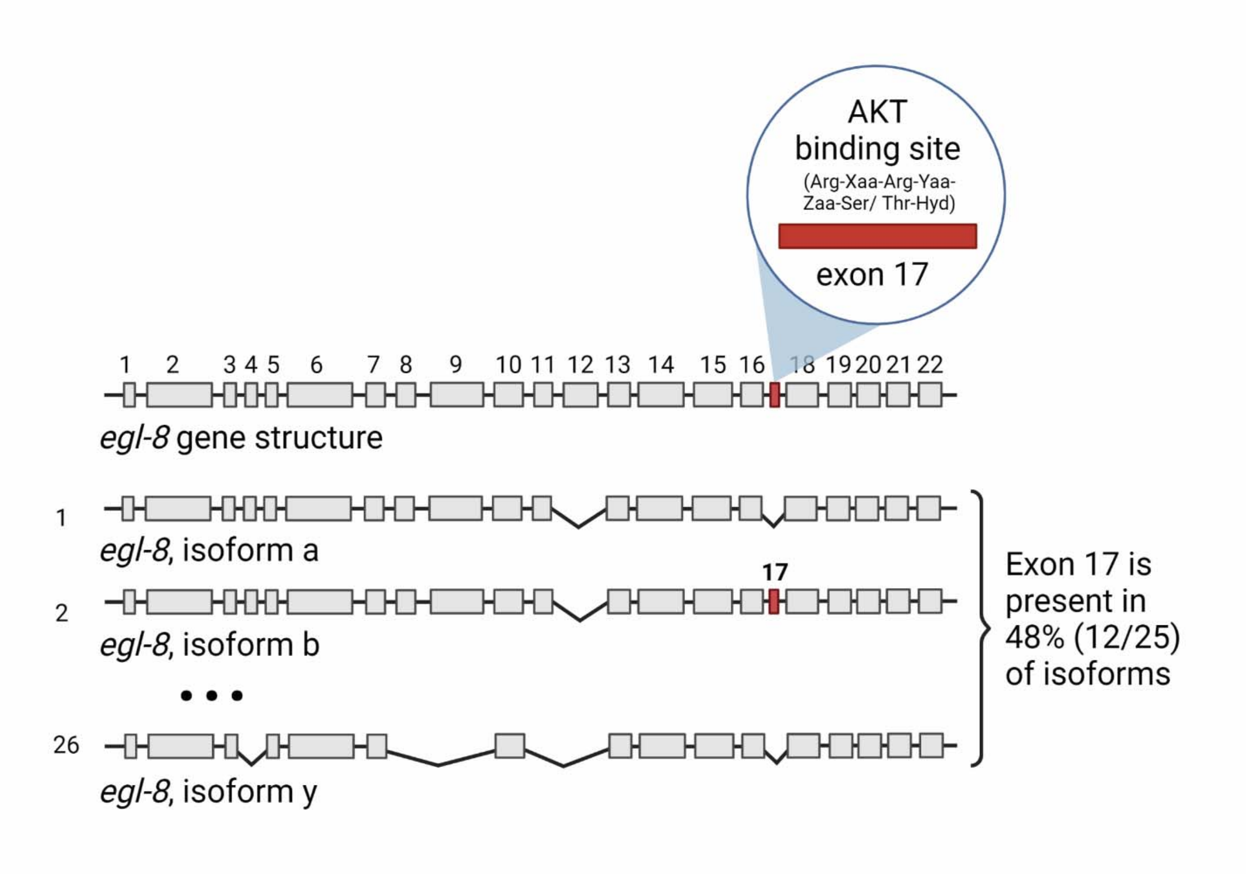


**Supplemental Figure 5** Location of the AKT binding site within *egl-8.* The AKT binding motif is present exclusively in exon 17, which is included in 12 of 25 isoforms. Binding motif as per Alessi et al. (1996).

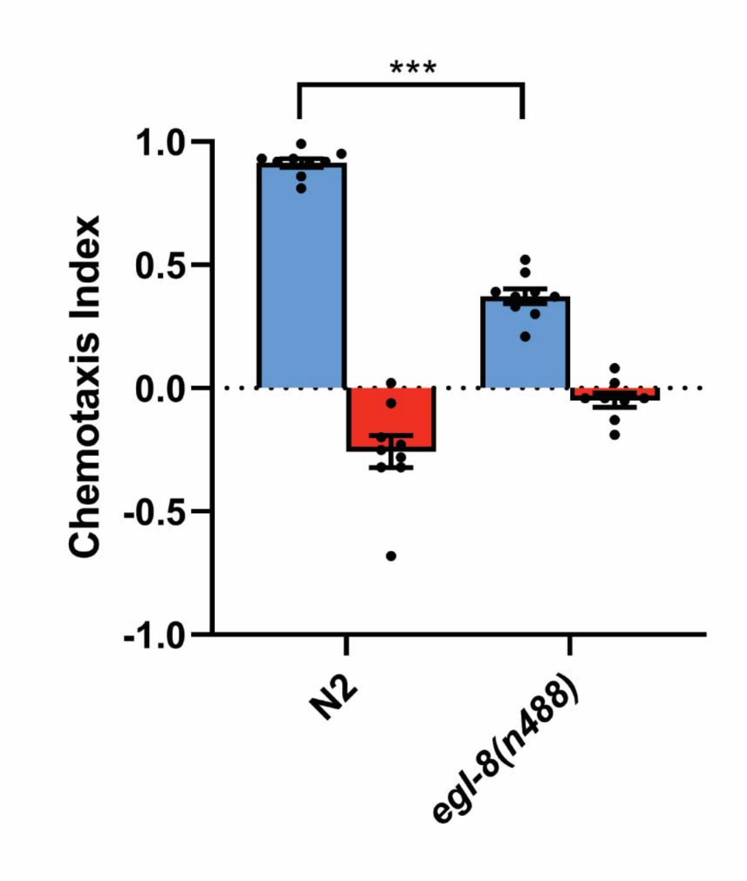


**Supplemental Figure 6** Chemotaxis of wild type animals grown at 20 °C for 52 hours after release from L1 arrest and *egl-8(n488)* animals grown at 25 °C for 52 hours after release from L1 arrest, to a point of benzaldehyde after benzaldehyde/starvation training.
